# Property Enhancer – a data efficient multi-objective approach for functional antibody optimization

**DOI:** 10.1101/2025.10.08.681101

**Authors:** Nataša Tagasovska, Jan Ludwiczak, Andreas Loukas, Emily K. Makowski, Homa Mohammadi-Peyhani, Sai Pooja Mahajan, Karina Zadorozhny, Donald Lee, Allen Goodman, Joshua Yao-Yu Lin, Ryan Kelly, Isidro Hötzel, Jack Bevers, Tamica D’Souza, James T. Koerber, Wei-Ching Liang, Julien Lafrance-Vanasse, Yongmei Chen, Andrew M. Watkins, Henri Dwyer, Stephen Ra, Richard Bonneau, Kyunghyun Cho, Vladimir Gligorijević

**Affiliations:** Genentech; Roche

## Abstract

In-silico antibody lead optimization remains challenging due to scarce high-quality data, costly experimental validation, and the need to jointly optimize multiple developability properties. Discovery workflows often rely on high-throughput phage, ribosome or yeast display experiments, which yield large but noisy datasets; as leads emerge, strategies shift to low-throughput assays which are precise, yet unscalable. Deep-learning and language-model approaches are hindered by such limited, unreliable measurements. We introduce Property Enhancer (*PropEn*), a data-efficient framework for low-data, heterogeneous regimes that can simultaneously optimize multiple antibody properties. PropEn proposes a matching-based augmentation that expands the training data with sequence *pairs* differing by only a few mutations; within each pair the second sequence improves the target value, providing an implicit optimization signal. Extensive *in silico* and *in vitro* tests show 10–39× affinity gains across four targets and nine leads, and enable joint multi-property optimization, positioning PropEn as a scalable, general solution.

## 1 Main

The process of drug discovery (DD) is inherently data constrained [1]: with each target-specific project typically yielding small low-throughput dataset of experimentally characterized molecules, often representing out-of-distribution (OOD) sequences relative to the public chemical and biological databases used to train generative models or large language models (LLMs). Although artificial intelligence (AI) and machine learning (ML) have shown promise in addressing many facets of pharmaceutical research [2, 3], a robust, standardized pipeline that can reliably guide lead optimization across diverse targets and diverse properties remains lacking. Fundamental challenges—scarce high-quality data, high measurement noise, and the sparsity of functional regions in chemical and biological space—undermine the performance of deep learning (DL) models, which typically require large, high-fidelity datasets to generalize [4],[5].

Experimental strategies, from targeted mutagenesis and high-throughput mutation scanning to evolutionary algorithms [6], attempted to navigate these limitations but fall short due to the absence of accurate predictive models to prioritize candidates. Analogously, ML-driven optimization approaches, including generative models steered by differentiable surrogate functions, succumb to the same core constraints: data scarcity, limited precision, and an inability to generalize across projects. This deficiency pervades various generative frameworks—variational autoencoders (VAEs) [7], diffusion models, and LLM-based methods—which, although capable of proposing diverse molecules from extensive datasets [8], struggle to optimize properties for a new target-specific antigen, as we demonstrate in our Results (2) section. To address this gap, we propose a novel tool tailored for the lead optimization phase of every DD project, designed to generalize across targets modalities and workflows, using smaller target-specific datasets.

### Our Approach

Here, we introduce Property Enhancer (PropEn)—a guidance-free lead optimization framework designed to improve any molecular property of interest. PropEn is well suited for low-data regimes, requiring as few as a hundred training samples, which our *matching-based augmentation technique* [12] transforms into datasets of thousands or ordered pairs—enabling the training of more expressive models.

PropEn, Figure 1 a - c, operates by *constructing molecular pairs*, where each pair consists of two similar molecules (e.g., antibodies), with the second molecule exhibiting an improved property. Such a setup allows us to train a deep model using a simple reconstruction loss: given a molecule with a lower property value, PropEn learns to reconstruct its higher-performing counterpart. Since paired molecules are highly similar in both feature and property space, this training paradigm *preserves context (proposing in-distribution, viable candidates) while implicitly optimizing molecular designs*. PropEn achieves higher gains with significantly smaller training sets compared to baseline diffusion (LAMBO, VDM) and language models (Efficient Evolution). The method relies on a much lower model capacity, denoted by the marker size in Figure 1 d. *In vitro* results (Figure 1 e). Collectively, these results show that PropEn, in the case of competing objectives such as binding affinity and expression yield, proposes designs closer to the Pareto front compared to state-of-the-art baselines.

**Fig. 1:**
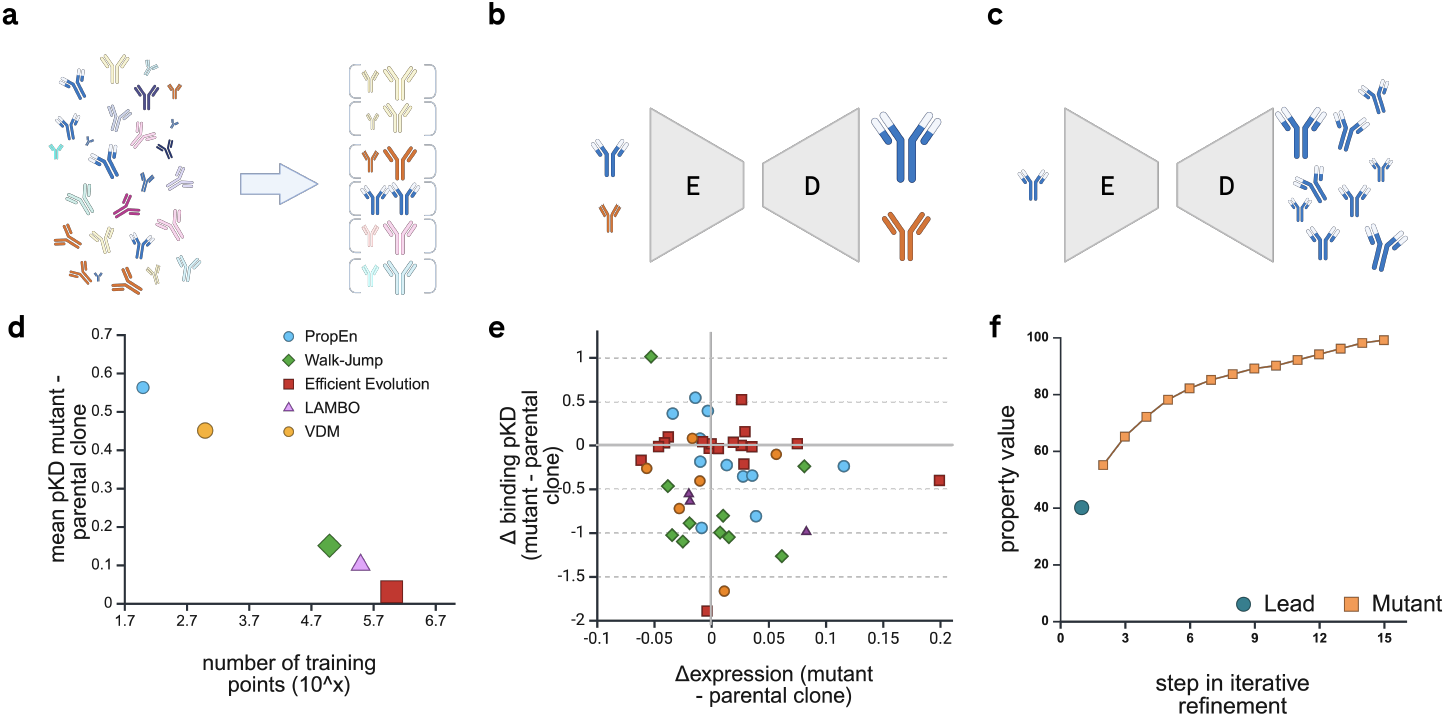
Property Enhancer (PropEn): training workflow and comparison to baselines. **a**, - as a first step, each antibody is matched to the most similar one in the training data, which exhibits a higher value for the desired property (ex. binding affinity). **b**, - at step two, a deep encoder-decoder model is trained with the matched dataset, i.e. implicit guidance, learning to map antibodies with lower to higher property values. **c**, - at step three, once trained, PropEn generates thousands of optimized mutants given a parental clone. **d**, PropEn achieves higher affinity gains with significantly smaller training sets compared to baseline diffusion (LAMBO [9], VDM [10]) and language models (Efficient Evolution [11]), at the same time, relying on a much lower model capacity, denoted as the marker size. **e**, - [*in vitro* results] PropEn simultaneously enhances binding affinity and expression across nine lead molecules from four distinct antigen targets. **f**, - sampled mutants can be further refined by iteratively re-feeding outputs to inputs into the encoder-decoder module.

Our PropEn framework offers three key advantages:

- **Eliminating the need for surrogate models** – Instead of relying on an external predictive model for guidance, PropEn directly learns a mapping from lower to higher property values, sidestepping one of the major bottlenecks in ML-driven molecular optimization.
- **Ensuring functional relevance** – By restricting training to known viable molecular space, PropEn generates outputs that remain within biologically and chemically relevant regions.
- **Simultaneous multi-property optimization** - By optimizing a multivariate ranking score, we jointly optimize antibody candidates such that they are close to the Pareto front.

Finally, PropEn’s encoder-decoder architecture enables iterative design refinement: antibodies can be iteratively refined by the same model, producing an optimization trajectory that incrementally improves the lead candidate, Figure 1 - f. As we demonstrate *vide infra*, this approach outperforms traditional generative models in producing *highquality, functionally relevant molecular designs across multiple property optimization tasks*.

## 2 Results

### 2.1 PropEn generates *in-vitro* functional antibodies across multiple targets

We validate PropEn through in-vitro experiments across four distinct targets (IL-6, HER2, EGFR and OSM) and eleven seed molecules in total. The goal was to enhance affinity maturation while preserving or improving expression yield. Surface plasmon resonance (SPR) [13] experiments were conducted to assess binding affinity. Details of experimental procedures and data collection are provided in the Supplementary Information.

PropEn successfully optimized lead molecules across all targets, achieving a 94.6% binding rate and 98% expression, outperforming VDM (86%), WJS (62.9%), and LAMBO (44.2%). This success stems from PropEn’s joint optimization of affinity and expression (see Figure 2 - a), that ensures that designs remain within a functionally relevant distribution. By exploring local sequence space, PropEn identifies novel, yet structurally viable mutations, allowing for improved molecular function while maintaining multiple developability properties (see Figure 2 - e).

**Fig. 2:**
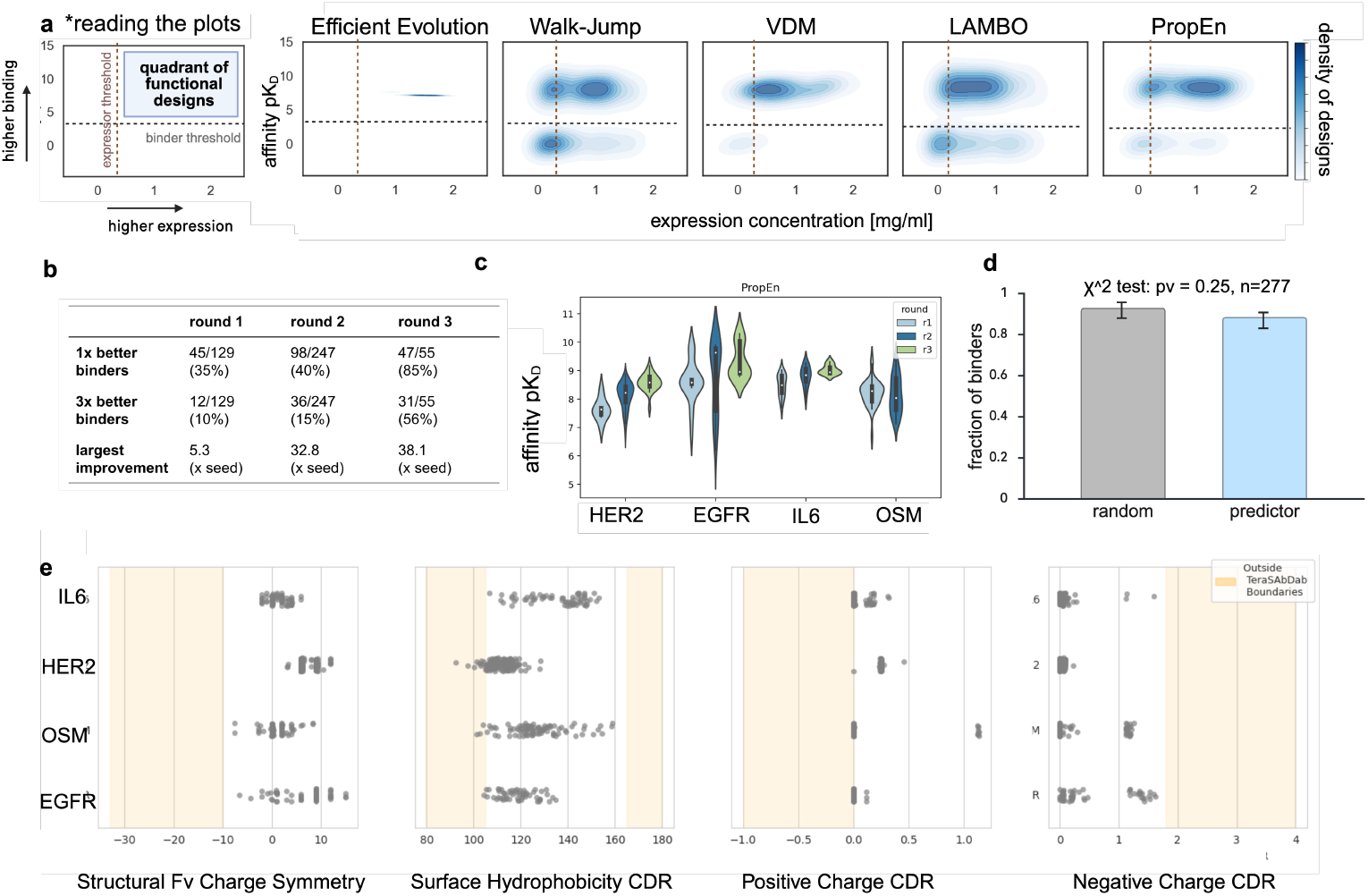
*In-vitro* results from 11 seed molecules and 4 distinct antigen targets. **a**, - Comparison of all baselines wrt expression (yield) and binding affinity. The first panel illustrates that a candidate design is considered functional if it surpasses the desired affinity and expression thresholds. Affinity pKD *<* 5 (defined as pKD = *−* log_10_(KD), where KD is the dissociation constant) is considered a non-binder. The distribution of PropEn designs is most concentrated in the upper right corner high binding rate and high expression rate. **b and c**, by running multiple measurement rounds of lead optimization PropEn designs improve leads across diverse set of targets. **d**, A/B testing confirms that there is no significance in using a predictor for selecting designs for in vitro experiments. **e**, developability properties of PropEn designs; PropEn provides not only good binders, but additionally antibodies with realistic therapeutic profiles computed by Therapeutic Antibody Profiler (TAP), with 96% of the designed molecules being within the valid therapeutic ranges.

### 2.2 PropEn is more sample-efficient than state-of-the-art ML methods

We compare PropEn against leading ML-based baselines trained on significantly larger datasets. *EffEvo*, for example, was trained on *millions of protein sequences*, while *WJS* and *LAMBO* leveraged OAS and internal datasets, and *VDM* was trained on 10x NGS sequencing data. In contrast, the PropEn models for each target were trained on datasets of only a few hundred sequences. Despite this significant difference in training data availability, PropEn outperforms all baselines in affinity maturation (Figure 1 - d), highlighting its ability to derive high-quality designs with minimal training data. This demonstrates that PropEn is not only effective but also highly sample-efficient, making it a promising solution for data-constrained lead optimization tasks.

### 2.3 PropEn does not require external guidance

As PropEn was evaluated as part of a larger effort in a lab-in-the-loop pipeline [14], its designs were passed through various predictors for selection. However, to guarantee that PropEns’ efficacy does not depend on the filtering method, we include results from an A/B test that was conducted in the same pipeline. Submitted designs were split into two groups: A - used predicted affinity pKD, binding probability, and per method ranking and B - used random (shuffled) affinity predictions and only global selection. Figure 2 - d, shows that PropEn rates appear balanced across both groups - control (random subset) and global selection including various predictors and heuristics, with no statistically significant difference between the two.

As shown in Figure 2 - d, Group A has about 94.2% binders and Group B has about 95.2%. Although B appears slightly higher, the χ^2^ test *p* = 1.0 indicates there is effectively no statistically significant difference between the two percentages. In other words, these results suggest that any slight difference in the fraction of binders between A and B could easily be attributed to random variation, so neither group is performing meaningfully better in that sense. This again shows that PropEn does not require any external guidance to achieve high-quality designs, because the guidance is implicitly built into the model.

### 2.4 PropEn discovers and enriches rare, meaningful mutations from complex datasets

In order to gain a more detailed insight into the nature of the mutations proposed by PropEn we analyzed the designs modifying the N032 anti-EGFR seed molecule. To this end, we have collated the results of the first design round where the available data around the seed molecule was scarce. We have found that essentially all designs improving the binding affinity of the seed molecule included a single G101D mutation in the CDRH3 region of the Fv - Figure 3 - a. This prompted us to investigate whether this particular mutation was present in the dataset used to train the method that contained 921 historical measurements of diverse naive- and affinity-matured clones binding EGFR antigen (at possibly various epitopes). No curation, clustering or other preprocessing was applied to this dataset before training. The G101D variant was indeed present in some of the clones yet its influence on the binding affinity was non-binary. The G101D mutation was present in both variants improving and worsening the affinity (Figure 3 - b) with no clear trend or enrichment. Interestingly though, the mutation was present mostly in clones that were characterized by larger distance to the seed molecule, including similarity in the CDRH3 region and in the close proximity to the seed (edit distance 0-4) it exhibited no effect on affinity (Figure 3 - b). This highlights that our strategy (cf. sequence matching section) and formulation of the method (cf. model architecture) allow to discover and enrich potentially non-obvious, property-improving mutations from complex, non-curated datasets. One potential drawback of this approach is that pairing between distant clones can introduce a large number of non-meaningful framework mutations [15] - e.g. best variant identified by PropEn for the N032 seed molecule had 7 edits, including 6 in the framework region (Figure 3 - c). Only the aforementioned G101D mutation had a direct and reproducible effect on affinity, most likely by introducing additional interactions with the EGFR antigen (exact crystallographic determination is pending). It’s worth noting though that excessive framework mutation can be circumvented both at the dataset preparation step (pairing) or as an additional post-processing step before experimental validation.

**Table 1.** % design candidates improving in-silico property by method. Higher is better, best numbers marked with bold.

|  | PropEn | Efficient Evolution | mPropEn | VDM |
| --- | --- | --- | --- | --- |
| Liabilities | **0.5** | 0.25 | 0.27 | 0.17 |
| Hydrophobicity | **0.56** | 0.34 | 0.33 | 0.47 |
| Charge - | **0.79** | 0.67 | 0.68 | 0.46 |
| Charge + | **0.65** | 0.41 | 0.29 | 0.51 |

**Fig. 3:**
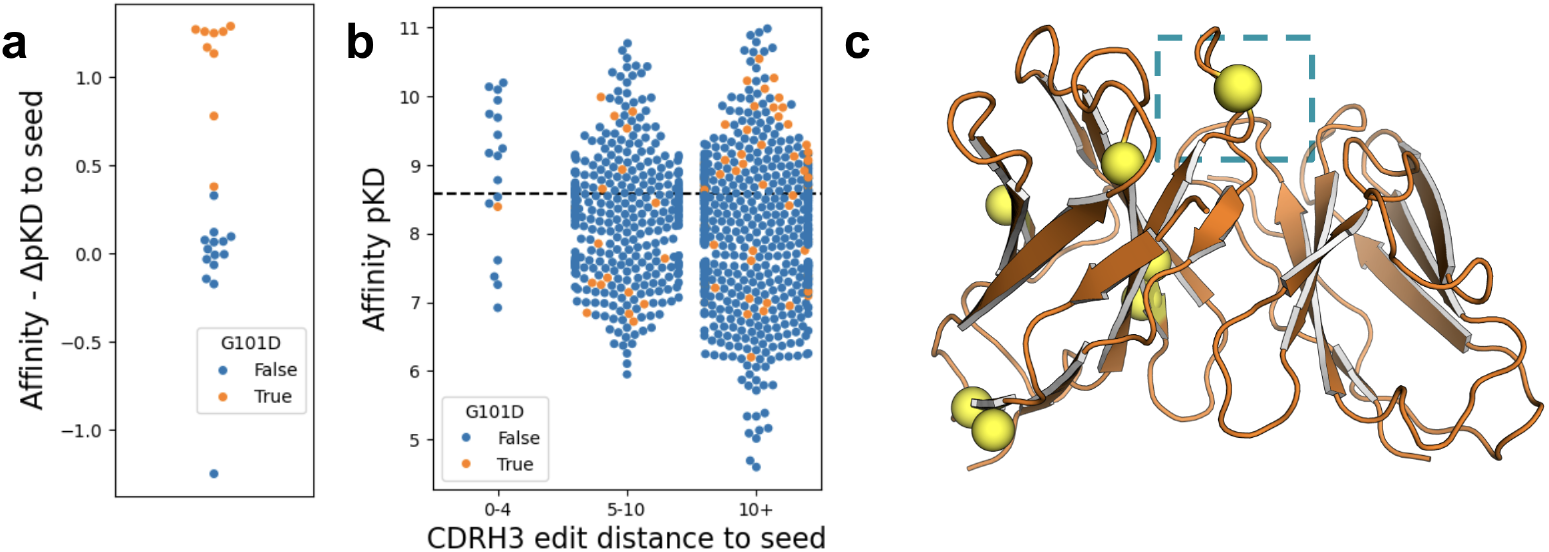
Discovery and enrichment of the affinity improving G101D mutation. **a**, Distribution of the binding affinities of PropEn designs modifying the N032-EGFR seed molecule. Model of the highest affinity design from this panel is subsequently shown at panel C. **b**, - Distribution of binding affinities in the training set binned by different similarities to the seed molecule in the CDRH3 region (annotated with the Aho definition, [16]). **c**, - Overview of the location of mutations in the best identified design mapped to the Ibex (CITE IBEX HERE) model of the design - modified positions are highlighted as yellow spheres. A single mutation modifying the CDRH3 region - G101D - is enframed in cyan.

### 2.5 PropEn optimizes diverse developability properties

We evaluate PropEn’s optimization capabilities using *in silico* experiments [17, 18], benchmarking its performance against state-of-the-art methods across multiple sequence- and structure-based properties of antibodies (see **Methods, Datasets**, for details). For each property, the objective is to **maximize the improvement per held-out test point** (Δ_property_(*parent*^*−*^ *mutant*)). A separate PropEn model is trained for each property, and the results—summarized in Figure 4 - d, —demonstrate consistent improvements across all properties. These results illustrate that PropEn achieves the highest Δ_property_, per-point improvement across all evaluation panels, highlighting its ability to generate diverse, yet functionally enhanced molecules. Moreover, in Table B1, we include the % of design candidates improving in-silico property by method. PropEn achieves highest rates from all of the baselines, providing between 50 - 80% improvement rate across all properties.

**Fig. 4:**
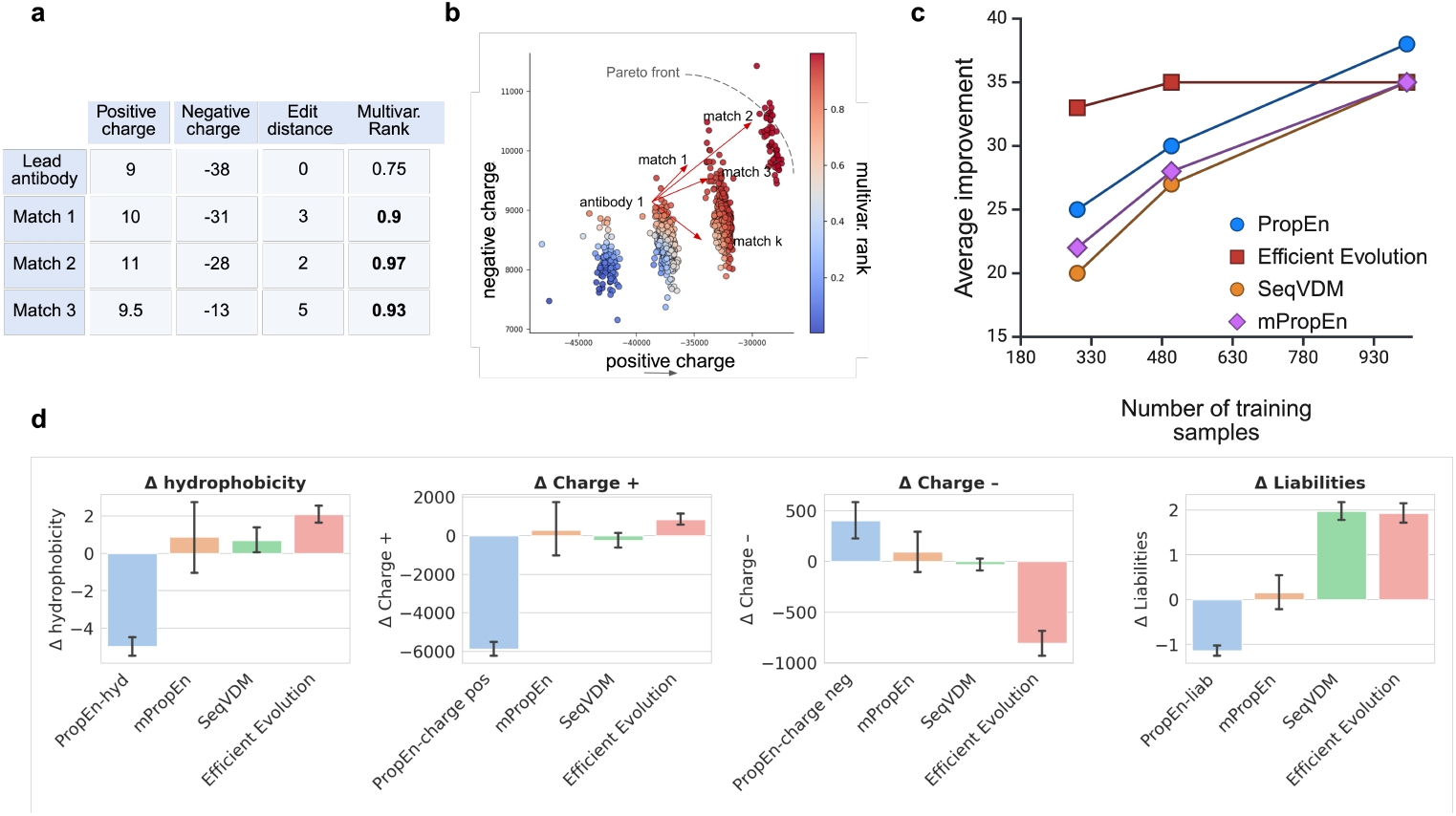
Multi-property optimization & in-silico evaluation. **a**, - An example of how a mutivariate ranking score is used in the matching. For a *Lead antibody*, each *Match* has one or more properties, as good as, or better than, the lead antibody’s resulting in a higher overall multivariate rank. **b**, - the multivariate rank score correctly orders antibodies, the ones close to the Pareto front exhibit higher scores (darker red). **c**, - sample efficiency of PropEn and mPropEn compared to baselines, **d**, - evaluating all methods wrt computational properties. Each method aims at reducing hydrophobicity, positive and negative charge and number of liabilities. Lower is better for all except “Charge”. PropEn consistently outperforms diffusion (VDM) and protein-language (Efficient Evolution) model baselines presented through delta in property value between each design and its corresponding lead antibody. mPropEn manages to keep balance among the competing objectives.

### 2.6 mPropEn enables simultaneous multi-property optimization

To evaluate PropEn’s ability to simultaneously optimize multiple properties, we introduce a *multivariate objective function* that ranks designs based on hydrophobicity, charge, and liability profiles. Thanks to multivariate scores computed BOtied [19], multi-objective PropEn, *mPropEn* enables asynchronous optimization— an approach not directly applicable to conventional differentiable optimization methods which require differentiable objectives such as [9]. Figure 4 a - d, show an overview of the integration of a MPO score within PropEn, enabling optimization of designs such that they approach the Pareto Front [20]. As shown in Figure 4 - e, multi-objective PropEn (mPropEn) achieves optimal trade-offs across all targeted properties simultaneously.

## 3 Discussion

We developed PropEn with multiple objectives: 1. a *target-agnostic* method that proposes a library of both diversified and optimized designs for a lead antibody; 2. *property-agnostic* method that can be applied to any property of interest; 3. *novelty in property values* - design antibodies with out of distribution, novel, property values 4. simultaneous *multi-property optimization* and finally 5. *no-guidance required*, removing the reliance on unstable, complicated design cycles that include both generative and discriminative modules. Our in-silico and in-vitro experiments confirm that PropEn does indeed achieve all of the above, primarily due to the pair matching procedure.

Conforming to the above design principles makes PropEn generalizable and efficient - only a small dataset of designs per (antigen) target is needed to succeed at producing binders in a single round of wet-lab experiments. Regarding data quantity and quality requirements, it is important to note that we had only couple of hundreds of points per target (statistics per target are listed in the Dataset section), collected from historical experiments across different teams in our organization; meaning there was heterogeneity, variety in the available antibodies per target as well. However, in the in-silico experiments and comparison to baselines, we used samples form a high-throughput mutagenesis dataset and the results were again in favor of PropEn.

The availability of in-silico, physics-based property scores, empowered us with an experimental setup to systematically evaluate our target and property agnosticism, as well as multi-property optimization features of mPropEn. Our in-silico experiments show the advantage of PropEn, in terms of absolute property improvement (largest delta between seeds and designs) as well as controlled generation. Language Model baselines have shown strength in proposing valid designs in close proximity to the leads, however due to lack of guidance, they do not improve the antibody with respect to the property of interest (Figure 2 a - efficient evolution does not provide any gain in binding), and WJS exhibits similar behavior. Guided baselines such as LAMBO are able to achieve high gains in binding by exploiting exploration, however, at the cost of producing a number of non-valid (poorly expressed) molecules. On the other hand, PropEn learns only “fateful” mutations by keeping the pairs in a matched batch close to each-other in all regards, except for the positions that lead to better property values. Learning these small but precise neighborhoods, results in proposing valid molecules with high certainty (*>* 90%).

## 4 Methods

### Overview of PropEn

PropEn is a matched-pair encoder-decoder framework that converts modest volumes of measured sequence–property data into a generative optimiser (Fig. C3). Unlike conventional gradient-guided pipelines—which combine an unconditional generator with a separately trained surrogate predictor and are therefore vulnerable to adversarial false positives[21, 22]—PropEn integrates the “improve” signal directly into the generative model via *matching*, eliminating the need for an explicit property predictor during optimisation.

### Matching strategy

For a dataset *Ƶ* = {(*x, y*)} of antibody sequences *x* ∈ {1, …, 20}^*L*^ (*L ≈* 250) and associated scalar properties *y*, we construct a set of *directed pairs*

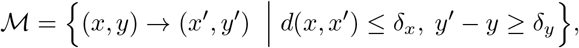

where *d* is the Levenshtein distance [23, 24], *δ*_*x*_ is the maximum allowed mutation radius (default 5) and *δ*_*y*_ is the minimum property increment (default 0.5 s.d. of the training distribution). See (Figure 5) **a**, for conceptual summary of matching. This procedure is inspired by nearest-neighbor matching in causal inference [25] and yields balanced “treatment” (higher property) and “control” (lower property) groups that differ only by local edits. In sparse optimisation landscapes, the hyperparameters *δ*_*x*_ and *δ*_*y*_ let practitioners trade off locality (exploitation) against coverage (exploration).

**Fig. 5:**
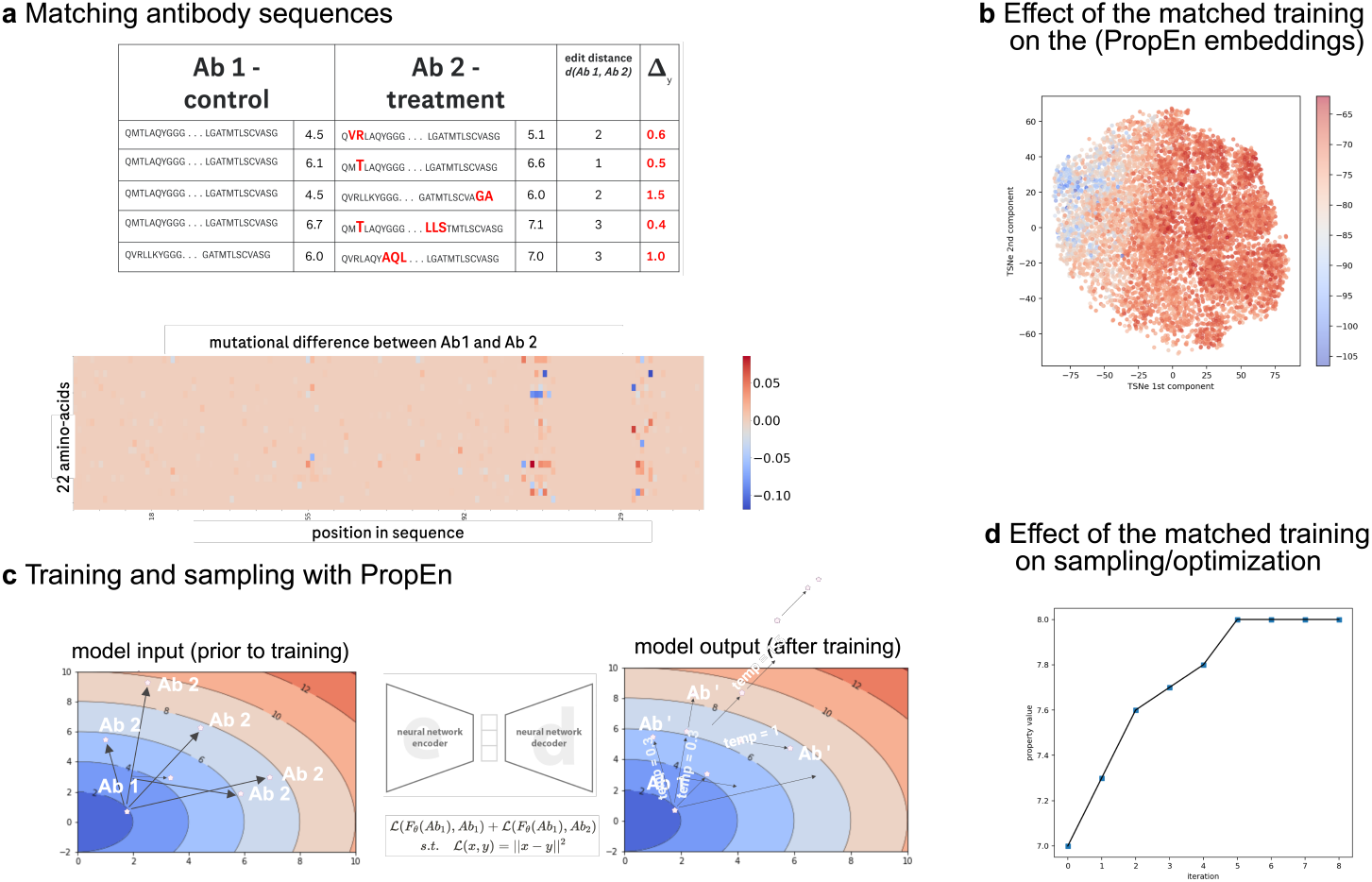
Method - property enhancement via matched training. **a** -, antibodies are paired by sequence similarity (low edit distance) into control (lower property) and treatment (higher property) groups; **b** -, these pairs train PropEn’s paired-loss to model a continuous property landscape and propose stepwise antibody improvements. **c** -, t-SNE of control (lower property) group antibodies, colored by integral surface hydrophobicity, reveals a learned monotonic gradient. **d** -, Iterative sampling along this implicit gradient produces successive property enhancements.

### Model architecture and training

PropEn comprises a three-block 1-D ResNet encoder (3.4M parameters) and a tied-weights decoder. One-hot AHo-aligned sequences, [16], are projected to a 128-dimensional latent space; ReLU activations and layer normalisation are used throughout. Given a matched pair (*x → x*^*′*^)*∈* ℳ we minimise the paired reconstruction objective

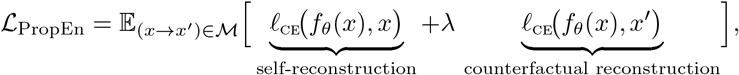

where *ℓ*_ce_ is position-wise cross-entropy and *λ*=1. The second term orders the latent manifold along the property gradient, a behavior we verify by visualizing t-SNE embeddings coloured by *y* (Figure 5). Training uses Adam (*β*_1_=0.9, *β*_2_=0.999), learning rate 1×10^*−*4^, batch size 128.

### Sequence generation

At inference time PropEn accepts a seed antibody *x*^(0)^ and offers two complementary generation modes.

#### 1. Iterative optimisation (default)

We repeatedly decode and re-encode, *x*^(*t*+1)^ = *f*_*θ*_ (*x*^(*t*)^), until *d* (*x*^(*t*+1)^, *x*^(*t*)^) = 0 or a user-defined edit-budget is exhausted. Property values typically plateau within five iterations (Figure 5, **d**).

#### 2. Stochastic sampling

The pre-softmax logits at each position define a categorical distribution; drawing from this distribution at temperature *τ∈* [0.1, 1.5] yields diverse variants in the vicinity of *x*^(0)^. Samples may optionally be refined with one or two further optimisation steps (hybrid mode).

### Multi-property extension - mPropEn

Drug-candidates must satisfy multiple, often competing, developability criteria [26]. We extend matching to the vector case by replacing *y* with a multivariate rank score *s* = BOtied(*y*_1_, …, *y*_*m*_)[19] and requiring *s*^′^− *s* ≥ δ_*s*_. The resulting *m*PropEn preserves Pareto optimality both in-silico Table B1 and in-vitro Figure 2.

## Baselines

Since recent progress in protein design is dominated by large-scale language and diffusion models, we benchmark PropEn against four state-of-the-art baselines, retrained on the same data when permitted by the original licenses.

EffEvo [11] Effevo leverages general protein language model (ESM-2) to suggest evolutionarily plausible antibody mutations—without any antigen, specificity, or structural input. It was shown it efficiently affinity-matures human antibodies by screening just a few variants over two laboratory evolution rounds.

WJS [27] trains a separate score- and energy-based models on a smoothed discrete protein sequence manifold, uses Langevin MCMC to “walk” through noisy sequence space, then “jumps” back to exact sequences via one-step denoising—enabling fast, non-autoregressive generation of diverse, high-quality antibody proteins.

VDM [10] A latent-space diffusion model trained end-to-end on one-hot AHo representations, [16]. We include both unguided (VDM) and ILVR-guided[28] (VDM-G) variants.

LAMBO-2 [9] A hybrid masked-language / diffusion architecture, a gradient-based guidance method for discrete diffusion models that enables direct sequence-space protein design—further enhanced by LaMBO-2, a multi-objective Bayesian optimization with edit-based constraints.

For in-silico experiments, all baselines were tuned for best validation performance under a fixed compute budget. The in-vitro experiments were ran by the corresponding authors of each paper, optimized for their best performance as suggested in the referenced publications. It is worth noting that each of these models leverages different architectures and training datasets, resulting in diverse cost - efficiency tradeoffs. Notably, PropEn is the smallest capacity model at the same time requiring smallest amount of data Figure 1.

### 4.1 Datasets

Our experimental evaluation is split into in-silico and in-vitro set of experiments. In what follows, we present the details and choices for each.

#### In-silico developability properties

In recent years, the use of in silico tools is increasingly more common to predict antibody developability and quality attributes [29]. Often, these are based on sequences and/or structure-derived parameters to screen and funnel initial hits or candidates into more promising leads. Hence, we propose a benchmark setup, where we can evaluate PropEn and some of the baselines for various properties of interest, individually and simultaneously. Following the advice and suggestions of antibody engineering experts, we choose to evaluate the properties summarized in Table B1.

#### In-silico data

For in-silico experiments we turn to a subset of [30]’s yeast display library of a singlechain fragment variable [scFv] of the parental clones 4D5. 4D5 is derived from the therapeutic antibody Trastuzumab and binds to the human epidermal growth factor re-ceptor 2(HER2). In order to maximize the number of function-retaining mutations while probing residues likely involved in binding, [30] chose two complementarity determining region (CDR) loops per scFv for modification: the CDRH3 and CDRL3 in 4D5. This procedure involved nine amino acids for mutagenesis in each antibody. The screening of the yeast display mutagenesis libraries were done with cognate fluorescently labeled antigens and sorted by FACS to obtain multiple populations. The 4D5 library contains three populations based on HER2 antigen binding strength (high-, low-, and non-binding).

From the provided libraries, we select the 4D5 population with high binding strength on the HER2 antigen. To mimic a low data regime, we subset 10% of the binders, resulting in a total of 11 556 antibodies. For each sample in the mutagenesis library, we compute the six in-silico developability properties. We chose structure and sequence based properties to demonstrate that PropEn can be effective even though it only uses sequence data. Each property is computed with a Python implementation of the procedures described in the the corresponding citations as listed in Table B1. The structures were obtained with Equifold [31], an end-to-end differentiable, SE(3)-equivariant, all-atom protein structure prediction model.

#### In-vitro data - binding affinity

Recombinant antibody expression and purification. Variable heavy- and light-chain genes were synthesized and subcloned into mammalian expression vectors, then cotransfected into Expi293F cells following the protocol of [14]. Cultures were maintained at 37 C with 8% CO and 125 rpm agitation. Five days post-transfection, supernatants were clarified by 4,000×g centrifugation and 0.22 *µm* filtration. Antibodies were captured on Protein A resin (GE Healthcare), eluted in 0.1 M glycine (pH 3.0) and immediately neutralized with 1 M Tris-HCl (pH 8.0), then buffer-exchanged into PBS (pH 7.4) using PD-10 desalting columns and concentrated to 1 mg/mL (see [14] for full details).

Binding affinity measurements. Equilibrium dissociation constants (*K*_*D*_) were determined by surface plasmon resonance on a Biacore T200 at 25 C. Target antigen was amine-coupled to a CM5 sensor chip at 500 RU. Antibody analytes were injected as a two-fold dilution series (0.1–100 nM) at 30 µL/min, with 180 s association and 600 s dissociation. Sensorgrams were reference-subtracted and globally fit to a 1:1 Langmuir model using the manufacturer’s software to derive *k*_*o*_*n, k*_*o*_*ff* and *K*_*D*_ (see [14] for full assay parameters).

## 5 Data Availability

The data for the in-silico experiments can be found in [30]. For the in-vitro experiments we refer to the original data source in [14].

## Declarations

### Funding

All work described in this paper was funded by Genentech Inc., South San Francisco, CA.

### Competing interests

All authors are or were employees of Genentech Inc. (a member of the Roche Group) or Roche, and may hold Roche stock or related interests.

### Ethics approval

All animal studies were performed in animal facilities accredited by the Association for Assessment and Accreditation of Laboratory Animal Care International. The procedures for animal studies were compliant under the Institutional Animal Care and Use Committee of the facility.

## 6 Authors’ contributions

Authors’ contributions K.C., V.G., and N.T. conceived of and designed the work. J.B., T.D., R.K., J.L.V., D.L., E.M., N.T., and Y.C. acquired and generated the data. J.B., T.D., V.G., R.K., J.L.V., D.L., A.L., J.L., S.P.M., E.M., H.M.P., and N.T. analyzed and interpreted the data. V.G., A.L., S.P.M., N.T., and A.M.W. created the methods and algorithms presented in this work. R.B., K.C., V.G., J.Y.Y.L., J.L., S.P.M., E.M., H.M.P., S.R., N.T., and A.M.W. wrote and edited the manuscript. R.B., K.C., V.G. and S.R. supervised the research., I.H. and J.T.K. supervised the execution of wet-lab experiments. H.D., A.G., S.P.M., N.T., and K.Z. performed computational experiments and developed code. All authors discussed the results and approved the final version of the manuscript.

## Appendix A Additional analysis on in-vitro data

### A.1 Comparing PropEn to NGS repertoire baseline

To contextualize PropEn, we compared it with a repertoire-mining baseline. For this baseline, we selected repertoire sequences whose pKD was comparable to or higher than the seed and generated variants by randomly recombining their observed mutations. Two caveats apply to this analysis: (i) the control set is *∼* sixfold larger than the PropEn set; and (ii) for PropEn we restricted analysis to seeds that were also included in the control experiment, which excludes two of PropEn’s best-performing seeds. These differences reflect the distinct provenance and aims of the experiments, but we include the comparison to provide perspective on how an ML-guided approach relates to a non-ML alternative.

First, the overall binding rate of PropEn is higher than the baseline, 91% and 86% respectively. Second, Figure A2 summarizes both the marginal (a, b) and joint (c) behavior of expression yield and affinity. The marginal plots show that PropEn designs are shifted toward higher values in both metrics: for yield, the distribution and median are higher (median shift +0.026 mg, 95% CI 0.016-0.038; Mann–Whitney p = 0.0049), and for affinity the distribution is likewise shifted upward (median shift +0.281 pKD units, 95% CI 0.212–0.353; p = 6.9× 10^*−*5^). Consistent with these marginals, the joint density in panel c reveals two modes—a shared low-yield/mid-affinity mode (0.05–0.10 mg; pKD *∼* 8–9) and a second lobe unique to PropEn at *∼* 0.17–0.22 mg and pKD *∼* 9.0–9.5—indicating enrichment of mid-yield, higher-affinity designs while still covering the overlap region observed for controls.

#### Statistical significance of the results

Nonparametric and resampling analyses corroborate the visual differences between methods. Mann–Whitney tests indicate that the distributions of both expression yield and affinity (pKD) differ between PropEn and controls (yield: *p* = 0.0049; pKD: *p* = 6.99×10^*−*5^). Bootstrap estimates (20,000 resamples) show that PropEn is shifted toward higher performance: for yield, the difference in means is +0.020 mg (95% CI 0.011 *−* 0.030) and in medians is +0.026 mg (95% CI 0.016–0.038); for affinity, the difference in means is +0.264 pKD units (95% CI 0.191 − 0.336) and in medians is +0.281 (95% CI 0.212*−* 0.353). Together, these results quantify the observed enrichment of PropEn designs toward higher yield and affinity relative to the control set in this dataset.

**Fig. A1:**
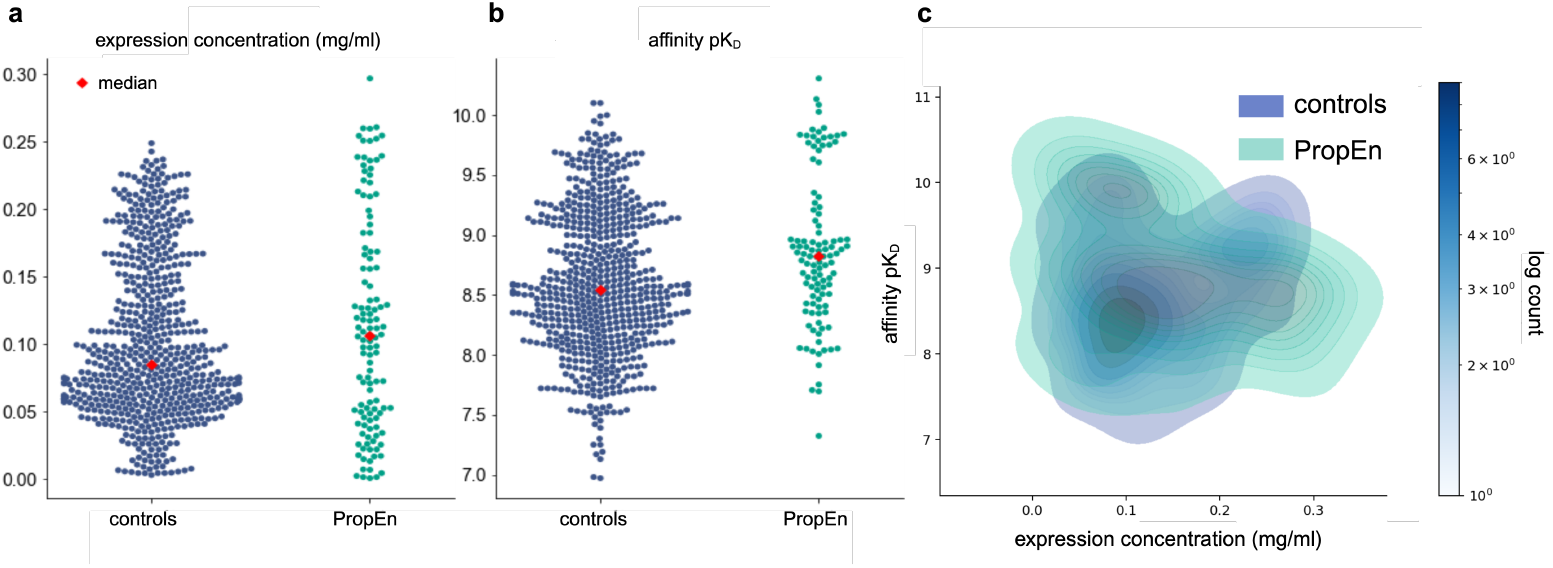
Marginal and joint distributions of expression yield and affinity for engineered designs. (a) Expression yield (mg) for controls (blue) and PropEn designs (orange), with medians indicated; PropEn is shifted toward higher yields (Mann–Whitney p = 0.0049; median shift +0.026 mg, 95% CI 0.016–0.038). (b) Affinity (pKD) for the same sets, also showing a shift toward higher affinity for PropEn (*p* = 6.910^*−*5^; median shift +0.281 pKD units, 95% CI 0.212–0.353). (c) Two-dimensional kernel-density estimates of yield versus affinity. Both sets share a low-yield/mid-affinity mode (0.05–0.10 mg, pKD *∼* 8–9); PropEn additionally shows a mid-yield/high-affinity lobe (*∼* 0.17–0.22 mg, pKD *∼* 9.0–9.5), consistent with the marginal shifts in (a) and (b) and indicating enrichment of mid-yield, higher-affinity designs.

### A.2 Comparing PropEn to human expert picks from NGS repertoire data

The results in Figure A2, across all three targets, PropEn reliably delivers high-affinity binders, with affinity distributions that are similar to or higher than those obtained from human-expert NGS selections. Expression is likewise competitive or improved: PropEn closely matches expert yields for EGFR and shows a clear upward shift for IL6; for OSM, PropEn spans a wider range that includes a subset of strong expressors, indicating room for straightforward tuning of expression objectives without sacrificing binding. Overall, these results demonstrate that PropEn can match or surpass expert curation on affinity while maintaining favorable expression, highlighting the method’s potential to scale design across targets with minimal trade-offs.

## Appendix B Details on in-silico experiments

## Appendix C Additional figures

**Fig. A2:**
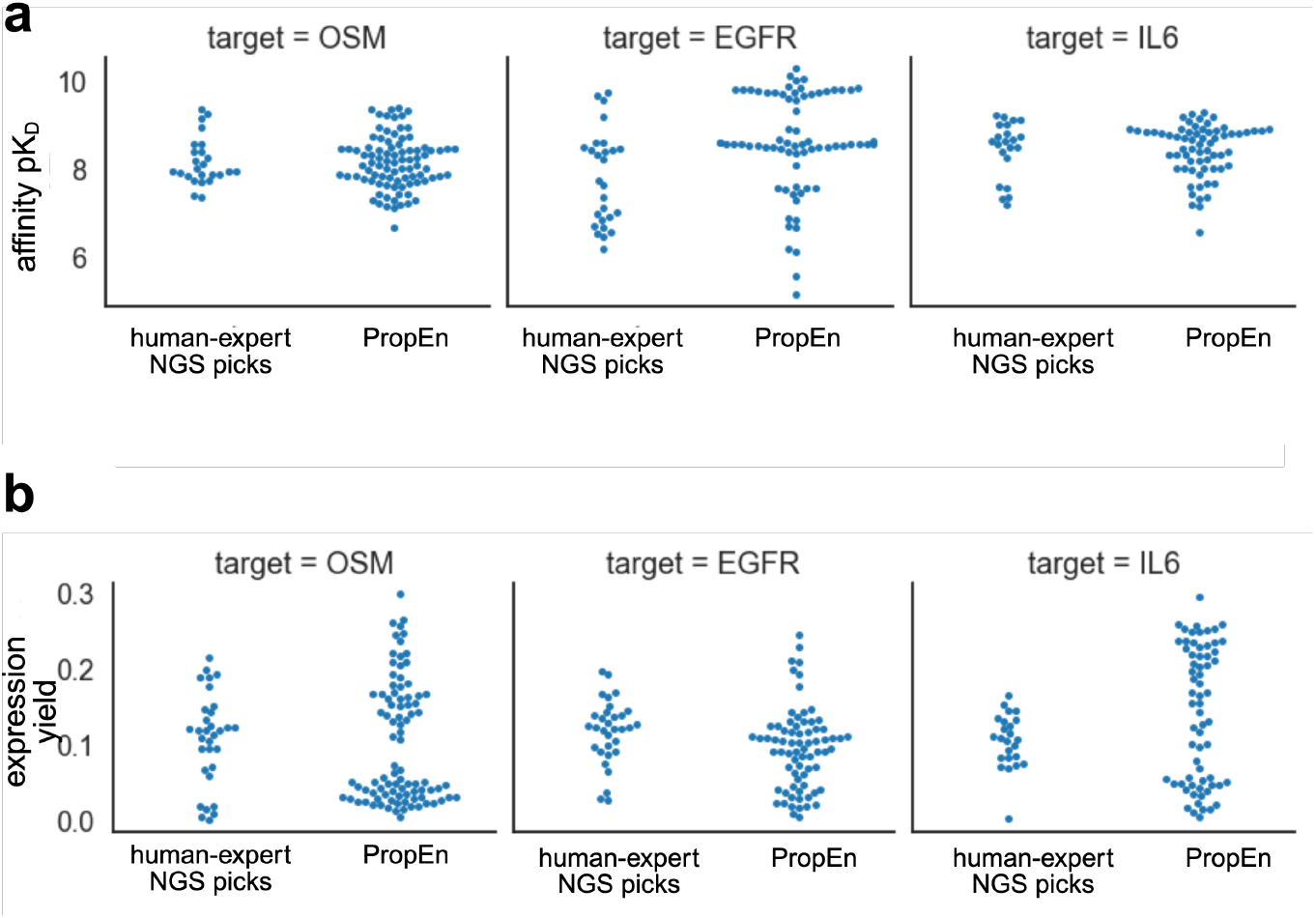
Distribution of binding affinity and expression for human-expert versus PropEn designs across three targets. a, Binding affinity pKD; higher values indicate tighter binding) for constructs selected by a human-expert NGS workflow and by PropEn for OSM, EGFR, and IL6. b, Expression yield for the same sets of constructs. Each dot is a single sequence; swarm jitter reflects local point density within each method group. Axes are independent across panels.

**Table B1.** Overview of in silico properties of interest.

| Property | Description | Objective | Dimension | Type |
| --- | --- | --- | --- | --- |
| hydrophobicity | Tendency of an analyte to prefer a non-aqueous environment. It may affect pharmacokinetics, efficacy, dose intervals, and route of administration [32]. | minimize | 1 | structure |
| positive & negative charge | Surface patch integral; the whole Fab is taken into account [33]. | minimize | 1 | structure |
| liabilities | Liability motifs are collected from the literature [34]. The goal is to minimize their count. Prioritize them by importance, with 1 being most important (e.g., W in CDRs) across the CDR vicinity. | minimize | 3 | sequence |
| multi-property score | Aggregates performance across all properties relative to a Pareto front. The Pareto-optimal solution has a score of 1.0, [19]. | maximize | 6 | sequence & structure |

**Fig. C3:**
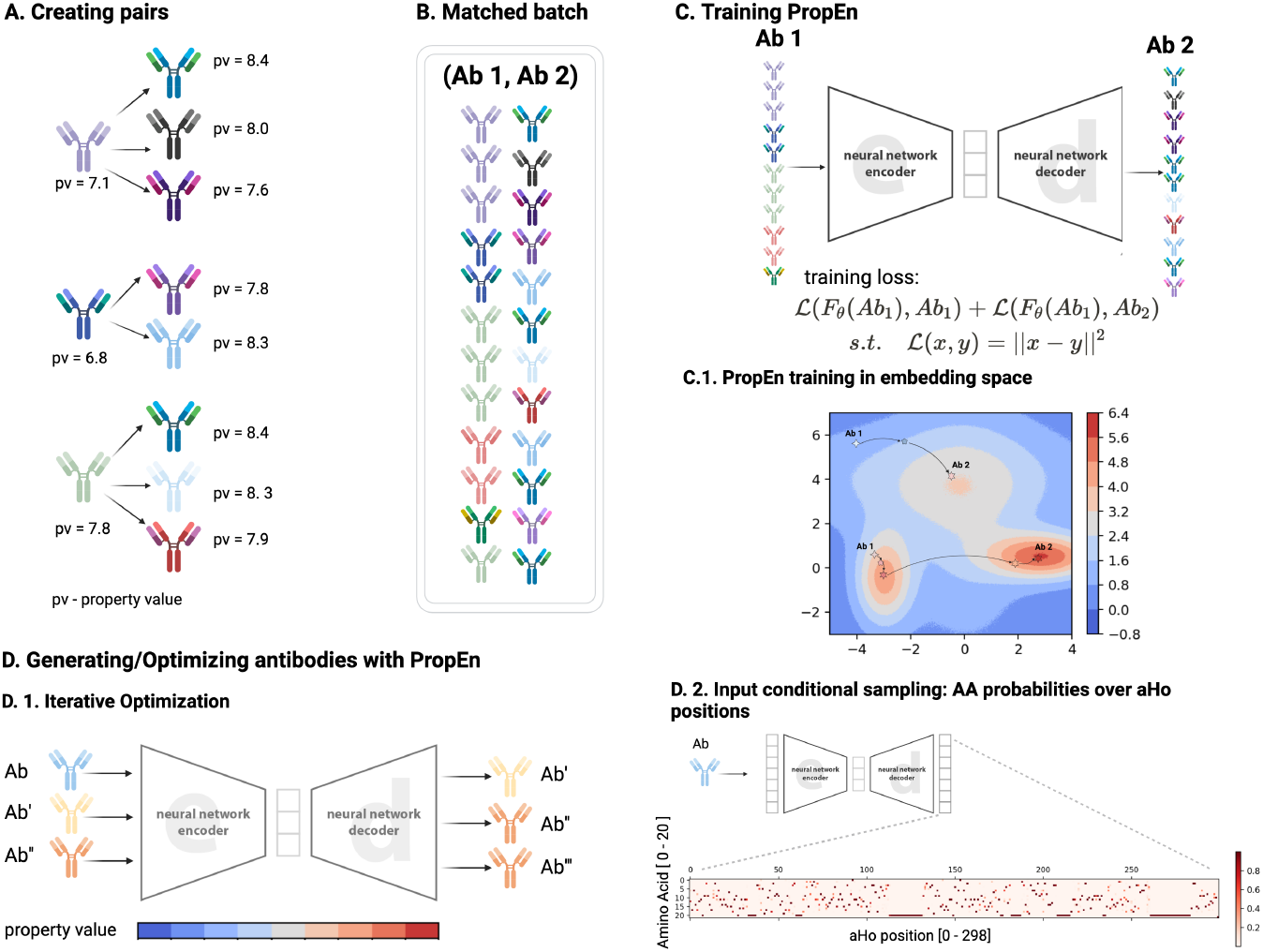
Property Enhancer (PropEn) for antibody lead optimisation. **a**, For every antibody in the training set we identify a *counterfactual neighbour* : a sequence that differs by at most *δ*_*x*_ point mutations yet exhibits a higher value of the scalar property of interest (Δ*y ≥ δ*_*y*_). **b**, These matched pairs form the minibatches used during training. **c**, PropEn is a lightweight ResNet auto-encoder that receives the lower–property sequence (Ab_1_) and is asked to reconstruct the higher–property partner (Ab_2_). **d**, After training, new designs can be obtained either (**d**_1_) by iterative re-feeding of the decoded output until convergence or (**d**_2_) by sampling from the final-layer categorical distribution at a user-defined temperature to trade off exploration and exploitation.

